# Time-Space Fourier κώ Filter for Motion Artifacts Compensation during Transcranial Fluorescence Brain Imaging

**DOI:** 10.1101/789370

**Authors:** Guillaume Molodij, Igor Meglinski, Anton Sdobnov, Yuri Kuznetsov, Alon Harmelin, Vyacheslav Kalchenko

**Affiliations:** Weizmann Institute of Science, Department of Veterinary Resources, Rehovot, 76100, Israel; University of Oulu, Optoelectronics and Measurement Techniques Laboratory, Oulu, 90570 Finland; Interdisciplinary Laboratory of Biophotonics, National Research Tomsk State University, Tomsk 634050, Russia; Institute of Engineering Physics for Biomedicine (PhysBio), National Research Nuclear University (MEPhI), Moscow, 115409, Russia; Aston Institute of Materials Research, School of Engineering and Applied Science, Aston University, Birmingham, B4 7ET, UK; School of Life and Health Sciences, Aston University, Birmingham, B4 7ET, UK

**Keywords:** transcranial imaging, *in vivo* fluorescence imaging, non-invasive optical imaging, motion artifact removal

## Abstract

Intravital imaging of brain vasculature through the intact cranium *in vivo* is based on the evolution of the fluorescence intensity and provides an ability to characterize various physiological processes in the natural context of cellular resolution. The involuntary motions of the examined subjects often limit *in vivo* non-invasive functional optical imaging. Conventional imaging diagnostic modalities encounter serious difficulties in correction of artificial motions, associated with the rapid structural variations and fast high dynamics of the intensity values in the collected image sequences, when a common reference cannot be provided. In current report, we introduce an alternative solution that utilizes a Fourier Kappa-Omega filtering approach. We demonstrate that the proposed approach is effective for image stabilization of fast dynamic image sequences. The validation of the Fourier Kappa-Omega filtering was performed on the images obtaining during mouse transcranial brain imaging using fluorescent microscope as well as on the simulated sequences of images. The proposed technique can be used autonomously without supervision and assignation of a reference image.

## 1. Introduction

Over the last decades a considerable attention has been given to optical-based imaging microscopy modalities especially to the intravital microscopy [1]. To enhance the quality of microscopy images in visible and near infrared spectral range contrast materials are very often in use [2, 3]. Optical imaging has an ability to acquire data at high speeds; a feature that enables it to not only observe static distributions of contrast, but to probe and characterize dynamic events related to physiology [4–8], disease progression and acute interventions in real time [1]. For example *in vivo* intravital microscopy (IVM) has become the mainstream technology for life science and it is used in a various ways in order to extract quantitative information about essential parameters including cell location, cell motility, cell interactions or blood flow [1–8].

One of the challenges in all live organisms imaging studies is related to the compensation of motion artifacts. Those motion artifacts should be compensated using active control approach (e.g. mechanical stabilization or gated/triggered image acquisition schemes) [9] and if it not possible should be removed using various types of computation [10]. Most of the available computation solutions are dedicated to solve this problem and can do it with different levels of success [9,10]. Nevertheless there are concerns that available methods are not efficient in cases where motion artifacts are accompanied with strong structural changes inside the region of interest during image acquisition [4–7]. It is especially relevant to the recently introduced technique (by our group), the Trans cranial Optical Imaging (TOVI). TOVI is dedicated to use dynamic fluorescence as one of the contrast for imaging characterization and quantification of cerebral blood vessels through the intact cranium [4–7]. This fluorescence imaging technique is also characterized by fast dynamic changes of the forms and patterns over time related to the flow of the fluorescent material in the vessels. Nevertheless, natural movements made during the image recording produce distortions that are unique in each frame. These motions include rapid jerks or saccades, slower drifts, and high frequency tremors perturbing the temporal analysis.

Image registration is the process in which two or more images acquired at different times, from different sensors with different resolutions or dimensions, or from different viewpoints/perspectives are matched to one another [10, 11]. This matching is accomplished via a process in which all of the images in the data set are transformed and aligned into a shared coordinate system. It is often described as finding an explicit function that performs a mapping of a target image onto a source image [12]. Also, variations in the image has been classified by three major types for the better determination of appropriate transformation [11]. The first type is the variations due to the misalignment removed by a spatial transformation when registering. Since the application of an optimal transformation in this class will remove these distortions, the variations are called distortions. The second type of variations are also distortions but are not corrected by the registration transformation because that affects the intensity values. They may also be spatial, such as perspective distortions. Finally, the third type is called variations of interest, corresponding to the differences between the images that may be spatial or volumetric but are not removed by registration. The third type variations are due to scene changes for which it is not possible to use usual algorithms, because no easy-to-find landmark locations in the image can be found along the time sequence.

Our motivation is to introduce a relevant method that could be applicable for the stabilization of intra-vital images characterized by strong structural changes and significant motion artifacts during acquisition with variations of second and third types. The uncorrected distortions as well as the variations of interest, which together are called uncorrected variations, are not removed by registration since an exact match is not possible. The distinction between uncorrected distortions and variations of interest is important, especially in the case where both the distortions and the variations of interest are local, because the registration method must address the problem of removing as many of the distortions as possible while leaving the variations of interest intact [12].

The Fourier shift theorem was proposed for registration of translated images [13]. It computes the cross-power spectrum of the sensed and reference images and finds the location of the peak in its inverse. The Lagrange multiplier method uses the Fast Fourier Transform to determine phase shifts in 2D and 3D as indicated in Ref. [14]. The method shows robustness when time varying illumination disturbances occur [15]. The Fourier Mellin transforms are also used to register the images, which are not only translated, but also rotated and with a change of scale [16].

Herein we proposed an adaptive time-space Fourier method (the κώ Fourier filtering) that was initially designed to remove the effect of intensity oscillations of astronomical observations [17] and, to correct image motions at the limit of the detection due to the noise. We also demonstrate that this approach is able to correct the entire image sequences without supervision and assignment of a reference image. In Section 2, we present briefly the experimental setup to collect a fluorescent temporal sequence. The κώ method is described in Section 3. Current Image Quality Assessments (IQAs) are difficult to interpret in the particular context of fast dynamic intensity associated to versatile appearance of structures in the region of interest. Quantitative assessments using both artificially corrupted images and real microscopy [18] or using a dissimilarity index calculated between a reference image and each image in the sequence in order to detect the presence of artifacts have been recently proposed in intra-vital video microscopy [19]. We introduce a useful temporal code color-coding method to perform a quick evaluation of the image quality of the entire sequence in Section 4. The obtained fluorescent images collected during transcranial mouse brain imaging are shown in Section 5, whereas the discussion raised by the results of this study and other practical implications of the approach are represented in Section 6.

## 2. Experimental Setup and data collection

A standard fluorescent zoom microscope SZX12 RFL2 (Olympus, Japan) coupled with the CCD camera Pixelfly QE (PCO, Germany) are used for the image acquisition. Camera control and image acquisition are performed through CamWare software (PCO, Germany). A standard fluorescent illumination source was used, namely, a mercury short-arc discharge lamp. The local IACUC committee approved all experimental procedures. The Institutional Animal Care and Use Committee (IACUC) approved all animal procedures. Anesthesia was performed by intraperitoneal injection of ketamine (10 mg/kg) and xylazine (100 mg/kg) mixture, as described previously [4]. BALB/C mice were used in this pilot study in order to verify usefulness of the proposed method. As it was also previously described that following administration of general anesthetics, an initial cut was made, and the skin over the frontal, temporal, occipital and parietal regions was removed by blunt dissection [4]. The exposed area was constantly moistened with saline. The mouse was then placed under the microscope lens on a special mouse holder with a warming plate, which maintained constant body temperature of 37 °C and other vital signs. For fluorescence imaging, a dose of 0.01 mg fluorescein in a volume of 50 microliters was injected into the tail vein. Experiment duration was less than 1 hour, after which all animals were sacrificed by barbiturate overdose. After contrast material administration, 800 raw images (exposure time: 45 ms per frame) were acquired.

## 3. Time-space Fourier Filter κώ

The main objective of the proposed research is to remove as many of the distortions as possible while leaving the variations of interest intact in order to envision a global mapping of the brain vascularization. For instance, the temporal distribution of a fluorescent material through the vasculature is different for veins and arteries. Fig.1 (a) demonstrates color-coded image representing frame of the maximum fluorescence in each image sequence (first 300 images from the acquired stuck) pixel during mouse brain visualization. The reddish colors correspond to the arteries with the fastest contrast agent administration. The greenish colors correspond to the veins which has slower contrast agent administration ability comparing to the arteries. The bluish colors correspond to the vessels with the slowest contrast agent administration (e.g. sagittal sinus). Thus, it is clearly seen that different vessel types are not observed at the same time and are localized in different places, so that a reference image to apply usual registration do not match the entire sequence when trying to compensate the disturbing motions. There exist computational imaging algorithms that are able to register sets of images, even for super-resolution, but are inaccurate when dealing with strong structural changes and significant motion artifacts during the fast dynamic data collection [20–23].

**Figure 1.**
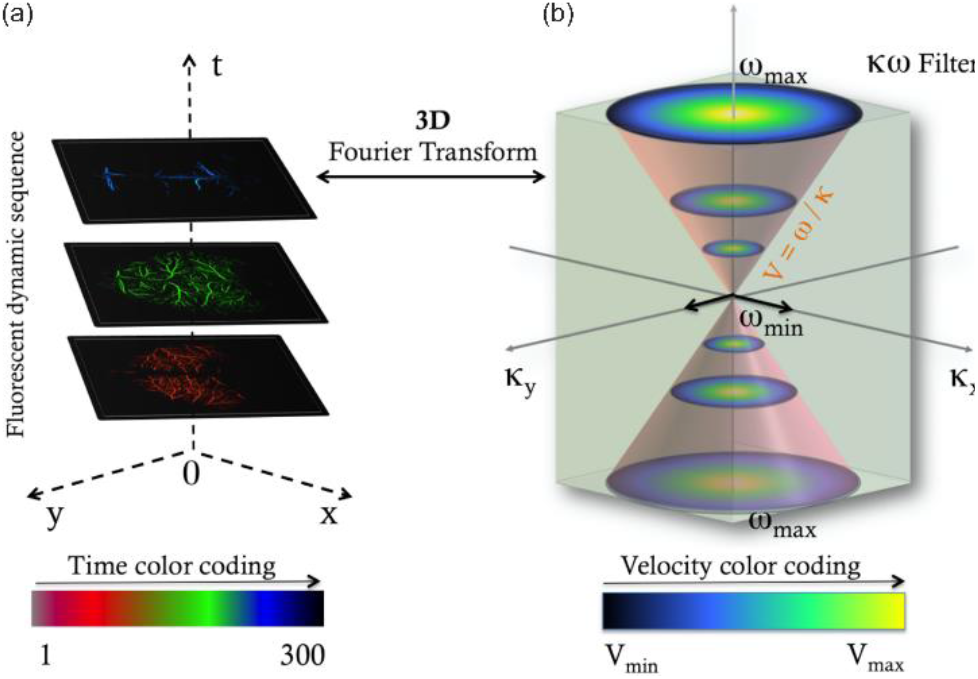
(a) Temporal color code (FIJI/Image J), represented by the color bar (Rainbow RGB code color) is applied along the frame sequence to distinguish the time-evolution structures (red for the arteries at the initial time, middle green for brain veins later and blue top for other veins (e.g. sagittal sinus) at the end). Anim8or-3D modeling freeware was used to display the figure. (b) Fourier time-space is represented taking into account the periodicity and symmetry properties of the Fourier transform; ώ_max_ appears twice while ώ_min_ is indicated at the center. The total sequence duration and the time sampling determine, respectively, ώ_max_ and ώ_min_. The pink cones delimit the removed frequencies of the Fourier space-time, defined by the velocity V = ω/κ. In the presented example v = 3.10^−3^ ms^−1^.

We propose to apply global motion compensation on the entire sequence using a time-space Fourier method, initially designed to remove the effect of intensity oscillations of astronomical observations [17,24]. The method consists in the determination of a three dimensional filter to adjust on the Fourier spectra of the raw fluorescent image sequence. Let us consider a single three-dimensional function of intensity *I*_*x*_, *I*_*y*_ for space, and time *t* that defines the fluorescent image sequences, with the notation *r*=*(x,y)*. The filtered time-space intensity is applied in the κώ space where κ and ώ are spatial and temporal frequencies. Let *Ĩ* = (κ, ώ) be the Fourier transform for space and time of the Intensity *I(r, t)*. In the spatio-temporal analysis assumption, the Fourier transform of the fluorescent signal *I(r, t)* consists of a projection of the temporal signal onto the orthogonal basis *exp*^−2*iπ*[ώ*t*]^ and the spatial signal onto the orthogonal basis *exp*^−2*iπ*[*r,k*]^,

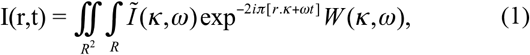

where the 3D time-space window filter *W*(κ, ω) is defined by

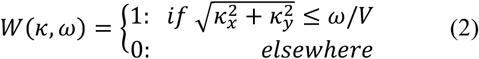

Fig.1 (b) shows a illustration of the space-time frequency representation of the Fourier analysis of the fluorescent signal. The use of a local spatiotemporal analysis allows the discrepancy of different regions in the Fourier space. For instance, the time-space filter of Eq.2 defines a cone of velocity *V* ≈ ω/κ that determines the velocity of the motions to discard. All Fourier components inside the κ-ώ volume are removed.

In Fig.2, Filters are displayed for different values of the ώ/κ ratio. The method acts as a selection of frequencies distributed in the 3D volume of the Fourier Space. In Fig.2, a linear relationship between space and time has been assumed to determine the values of the frequencies to remove in the κώ space. Another quadratic behavior might be also envisaged in the case of a diffusive law assumption.

**Figure 2.**
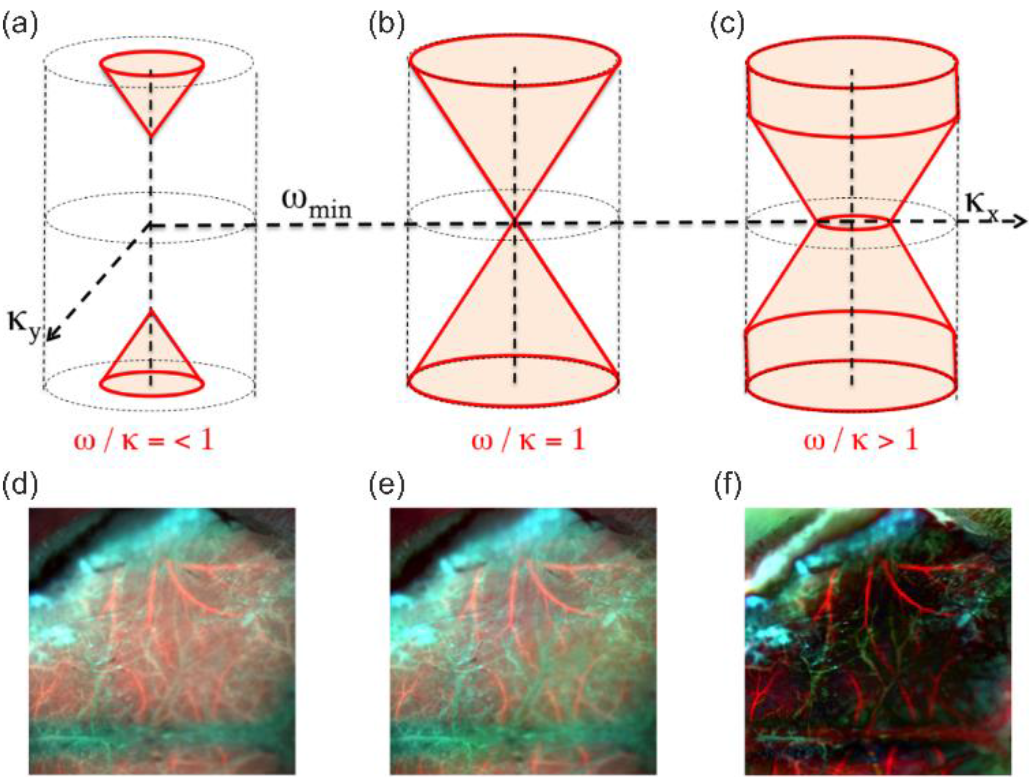
(a-c) The removed volume in the Fourier frequencies is indicated in pink. (d-f) Time color-coded images after the κώ filtering. From left to right the velocity criterion *V* = *ω/κ* is 0.5 (a,d), 1 (b,e) and 2 (c,f), respectively. The temporal color-coding (FIJI / Image J) is applied along the time sequence to distinguish the time-evolution structures from the image motions when stacking the 300 images.

As it was mentioned before, it is not possible to use reference image for motion artifacts compensation as far as fast structural changes such as arteries and veins are collected at different time and located ad different places. Fig.3 shows the result of the processing after applying a stationary signal detection uilizing reference image by comparison to the κώ method. The rigid registration method computes the cross-power spectrum of the sensed and reference images and finds the location of the peak in its inverse (implemented in the furnished Matlab code for comparison). The rigid registration method produces saccades due to the dissimilarities between the selected reference and the current image as indicated in Fig.3 (a) where is displayed the blurring coming from the missalignment after the registration.

**Figure 3.**
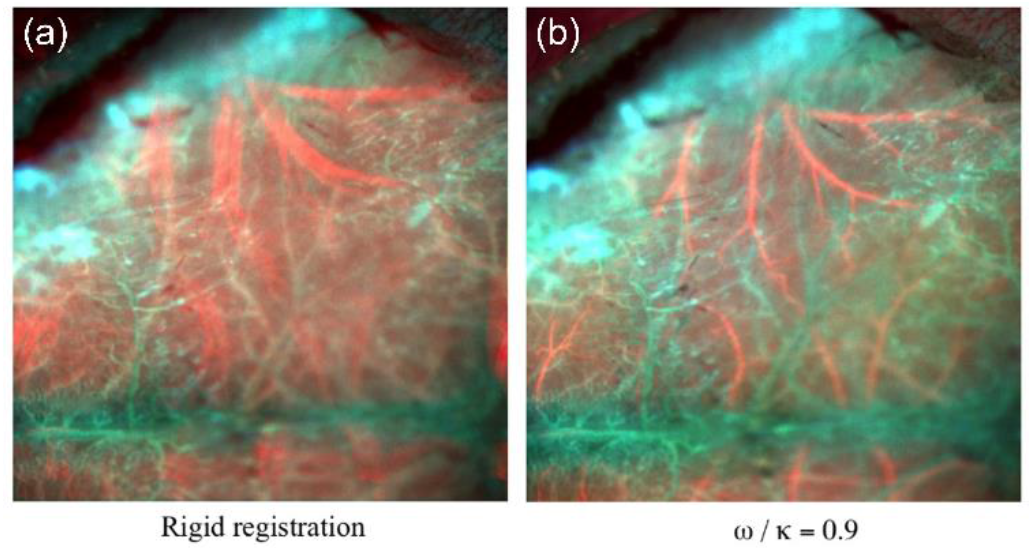
Comparison between a stationary detection approach (a) and the κώ method (b) when fast scene changes occurred. The temporal color-coding (FIJI / Image J) is applied along the time sequence to distinguish the time-evolution structures from the image motions when stacking the 300 images. On (a) the blurring of red structures (beginning of the sequence, for instance) is produced by saccades due to the registration missalignment.

## 4. Time color image distortion method

As a practical tool to obtain a quick look of the efficiency of the described method, we propose to use a temporal code color to overcome the difficulty due to the blending of the motion effects and the change of structures along the time. The main idea is to weight each image with a different color before the average in order to regroup structures appearing with the purpose to distinguish the change of structures from the motions. Color-coded maps are created using a multicolor merge approach of a full-unprocessed time sequences using the temporal color code plug-in in Fiji image analysis software (http://fiji.sc) [25]. Briefly, 300 frames are color coded and merged for a qualitative representation of image/time distortions. A “rainbow RGB” color scheme is chosen to correspond to the different level of structure appearance along the temporal sequence.

By comparison, we evaluated the image quality using methods such as the Similarity Image Assessment (SIA) [26], the entropy [27,28], and the sharpness [28] on the final obtained average image to compare with the time color image distortion method. This effort was firstly directed toward tuning quickly the κ/ώ parameter to find the optimum parameter. The lack of a common reference that fits the entire sequence leads us to adopt absolute assessments such as the entropy to estimate the quality of information retrieved or from the sharpness metric using the knowledge provided by adaptive optics in astronomy [28–30]. Sharpness assessment characterizes the features in the image while the entropy method indicates the level of information in each frame. An advantage for the entropy assessment is coming from its relative insensitivity to the intrinsic changes of the features. Optimum value of the filtering velocity parameter v=ώ/κ can be derived and eventually automated.

Moreover, we remarked that the κώ filter acts to remove the “background noise” due to the fluorescence scattering in the medium for large value of the velocity. As an example in Fig.4 (b,d), hidden structures emerge for recognition when applying the proposed filtering. Nevertheless, an accurate study of the image quality assessments to quantify the reliability of the background subtraction is beyond the scope of this paper. In regard only of the IQAs entropy scores and denoting that the κώ method depends on the temporal size of the sequence, we propose to adapt the κώ depending on the sequence time duration. In Fig.4 (a,c) the κώ filtering is not applied when ώ=ώ_min_ in order to keep the “background noise”. In the case of a long temporal sequence (as indicated in Fig.5), the processing leads to a similar result when determining the best filter, i.e., one obtains the same value of the optimum velocity from the IQAs.

**Figure 4.**
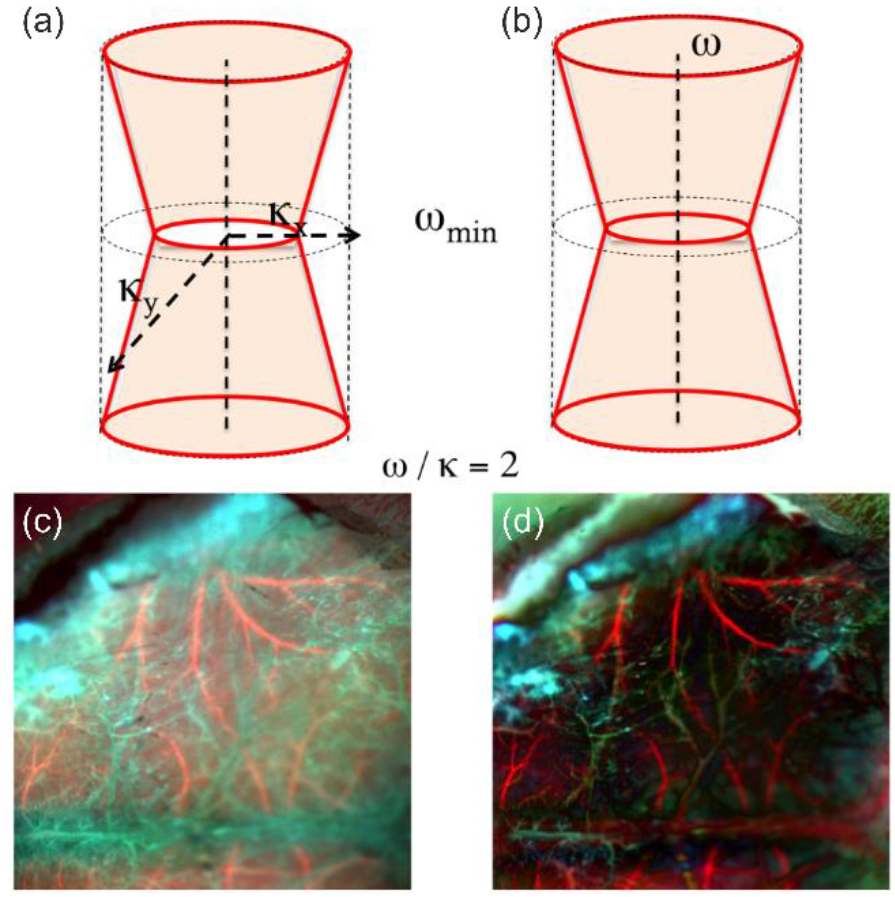
(a) The adaptation of the κώ filtering for short duration sequences. The frequencies at ώ=ώ_min_ remain intact. (b) Comparison to the method when the same velocity has been applied. (c,d) Corresponding time color image distortion renders.

**Figure 5.**
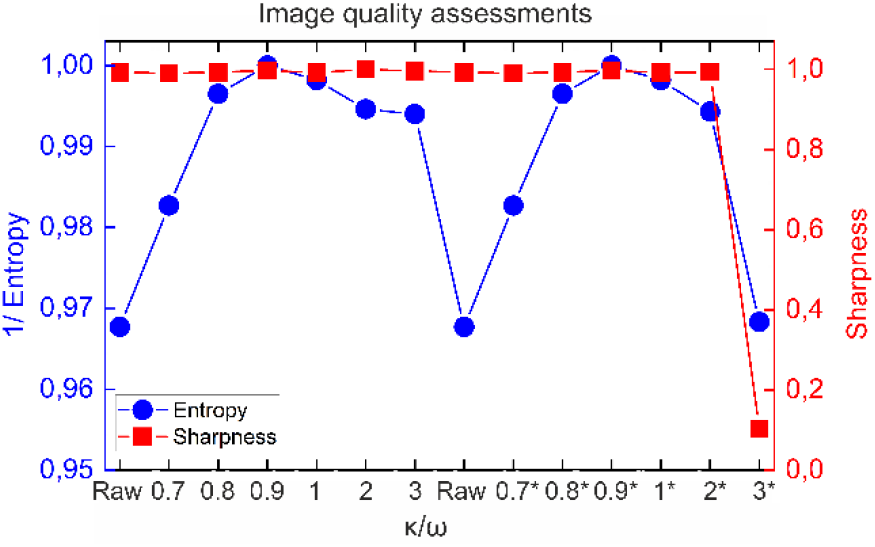
(IQAs (sharpness and entropy) are plotted for different values of the velocity parameter v=ώ/κ. Left part of the plot corresponds to the modified filter presented Fig.4 (top left), while the asterisk symbol mentioned on the velocity (x axis) correspond to the filtering indicated in Fig.4 (top right), i.e. for a progressive removal of the “background noise”.

In the case of short temporal sequences, applying a rigid registration eventually followed by the κώ would be more adapted, taking into account that it would be difficult for fast structural changes to appear during the collection. Fig.5 plots the IQAs values for a long temporal sequence case study. For sake of simplicity, the maximum score to display the values on the range from 0 to 1 normalizes all values. Image quality entropy assessment indicates a value of *v* = 0.9 as the optimum value for both types of filtering. Important to notice that sharpness of the long duration sequences is significantly decrease at the value of *v* = 3.

We verified the validity of the linear relationship assumption for the κώ filter by increasing the duration of the sequence (from 300 up to 800 frames). The corresponding optimum value of the velocity v=ώ/κ is then proportionally shifted.

## 5. Results

In the experiment related to Fig.3, the tissue was grossly immobilized, leaving only minor distortions in the field resulting from respiration and heartbeat. However, in some cases, a motion can actually shift the entire field of view. In this close-up image, the temporal sequence was made up 800 images and the growing factor scale was about 20 by comparison to the following experiment. There is an interest for an accurate compensation of jerks before any characterization or quantification of cerebral blood vessels through the entire intact cranium shown in Fig.6. This experiment was also an opportunity to apply the κώ method on an irregular data cube (x = 696 pix, y = 512 pix, t = 300 x 45 ms). Obviously, a better spatial resolution is obtained when applying the filtering. But important also, is the accurate recognition of the structures that appear at different moment and are grossly indicated by the color-coding. For instance, in Fig.6 (a), due to the involuntary motions, the discrepancy between veins structures (indicated in blue and green) at different instant becomes difficult. By comparison, in Fig.6 (b), one can expect to label the different functional vessels.

**Figure 6.**
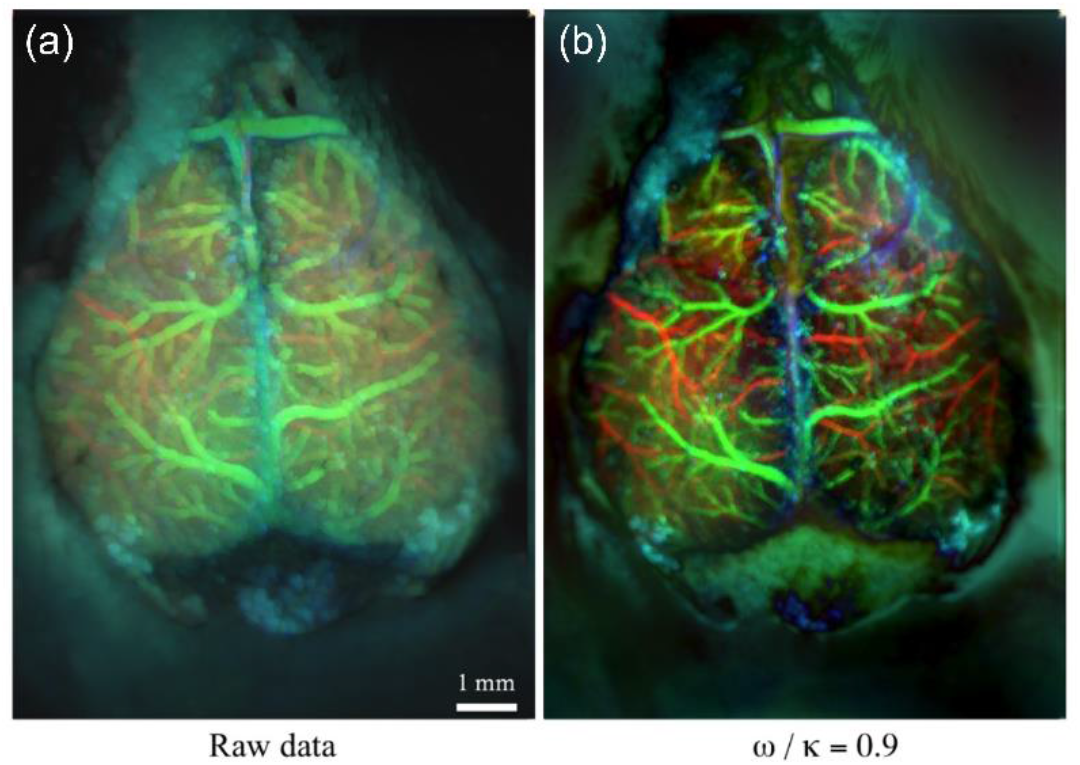
Dynamic fluorescence temporal sequence stacked over 300 images before and after processing. (a) The mean raw image sequence. (b) The same sequence, is displayed after applying the κώ method. Entropy IQA indicates a value of v = 0.9 as the best quote for the information quality (Fig.7). A temporal color coding plug-ins (FIJI/Image J) has been applied along the time sequence indicated by the color bar. The white bar corresponds to a size scale of 1 mm.

**Figure 7.**
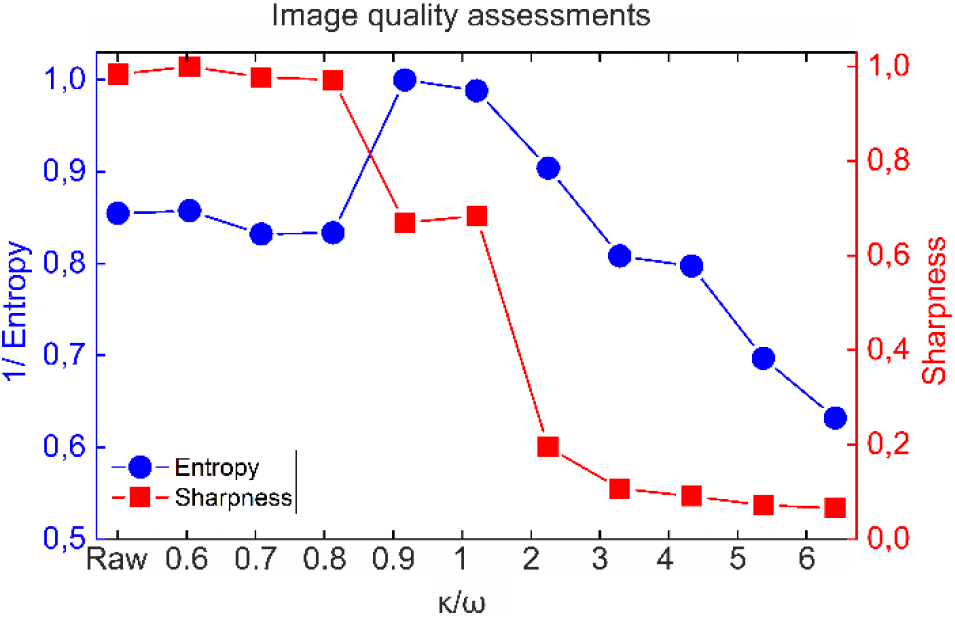
IQAs (sharpness and entropy) are plotted for different values of the velocity parameter v=ώ/κ. Sharpness assessment characterizes the features in the image while the entropy method indicates the level of information. Entropy and Sharpness are absolute IQAs (without calibration reference [29]).

In the κώ Fourier space mixed information is coming from image structures as well as dynamics. The velocity parameter ώ/κ delimits the Fourier space-time corresponding to the undesirable motions faster than the time dynamic scale of the structures. If the temporal sampling rate of the sequence is high enough, tuning the velocity parameter can separate both effects. Saccades from jerks are described by velocities larger than characteristic velocity of structure changes in the image, easy to extract.

Fig.8 displays the time color image distortion of the fluorescent sequence when increasing the value of the velocity. Image quality entropy assessment indicates a value of *v* = 0.9 as the optimum value for the registration (Fig.7). Important to notice that sharpness of the long duration sequences is significantly decrease for the value of *v* > 3.

**Figure 8.**
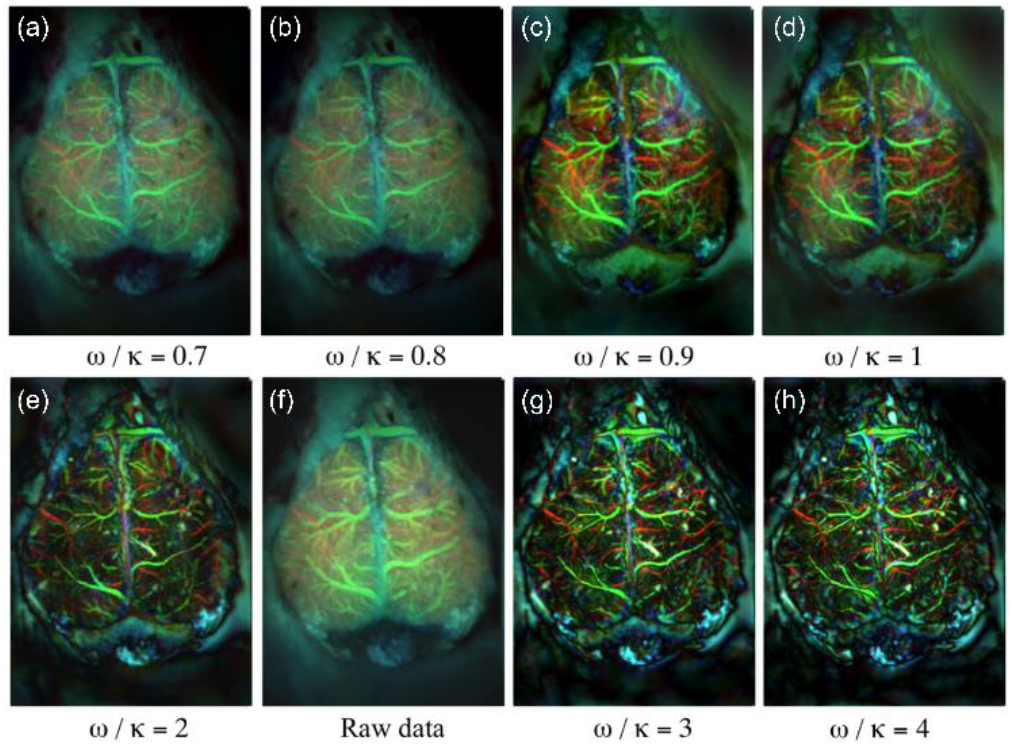
Comparison of the filtering efficiency on the same temporal sequence for values of the velocity ώ/ κ=0.7 (a), 0.8 (b), 0.9 (c), 1 (d), 2 (e), Raw data (f), 3 (g) and 4 (h). Image quality entropy assessment indicates a value of v = 0.9 as the optimum value for the registration (Fig.7). For this value, the filtering starts to reduce the background noise coming from the scattering due to fluorescence.

Nevertheless, increase the number of frequencies beyond the optimum to remove totally the jerks is accompanied by a loss of information in relation with the image structures. The high spatial resolution structures vanish and complex structures are mixed. Smaller looking vessel diameter for larger velocity parameter is due to some “erosion effect” related to the loose of corresponding frequencies. Images look sharp but we remark that details are vanishing. This is effect may explain the relative low scores of both entropy and sharpness IQAs for large value of the velocity in Fig.7. It is important to notice that both sharpness and entropy QIAs are applied directly on the average sequence image after κώ filtering without any correction of the dynamic of the signal that is incompatible with the definition of the entropy QIA (see the Matlab code).

## 6. Discussion

In this section, we discuss first issues related to the definition of the registration. The novelty here is that we apply the algorithm in a 3D space. Many of the problems arising from the projection of 3-space onto a 2D image may no be longer relevant [11]. From 2D studies, it has been accepted that the fundamental characteristic of any image registration technique is the type of spatial transformation or mapping used to properly overlay the images [11]. In this paper, our motivation was to introduce a relevant method to remove only some of the variations; the effects of illumination changes may be difficult to remove, or we are not interested in removing them, i.e., there may be changes that we would like to detect. So, the distinction between uncorrected distortions and variations of interest is important, especially in the case where both the distortions and the variations of interest are local. Because the registration method must address the problem of removing as many of the distortions as possible while leaving the variations of interest intact, it becomes necessary to distinguish between whether certain variations are global or local and whether the selected transformation is global or local. For example, images may have local variations, but a registration method may use a global transformation to align them. In the κώ method, the difficulty to display an explicit geometric transformation comes from the difficulty to distinguish the frequencies related to the global and local variations in the κώ space. Only an underlying geometric transformation law can be suggested.

In the purpose to determine the limitations and optimize the parameters, we applied the κώ processing on synthetic data. In the following simulation presented in Fig.9, the field of view is parsed with three-colored number of different sizes. Each color corresponds to a specific variation of the intensity indicated in Fig.9 (b). We simulated distortions such as the respiration and heartbeat together with random jerks. Plots of the generated motions are indicated in Fig.9 (c). The noise and the scattering effect are not generated. This simulation addresses the following purposes, in order to:

- Define spatial patterns for recognition (i.e., the three numbers) that are associated to three different time behavior (such as the brain vascularization),
- Produce saccades when applying a rigid registration (generating enough dissimilarities between the reference and the current image),
- Mix frequencies components close to the real case (such as the respiration and involuntary random jerks),
- Blend the information coming from the intensity variations and displacement motions in the Fourier space (that is not the case for a simple moving disk, for instance),
- Show the interest of the time color image distortion method.

**Figure 9.**
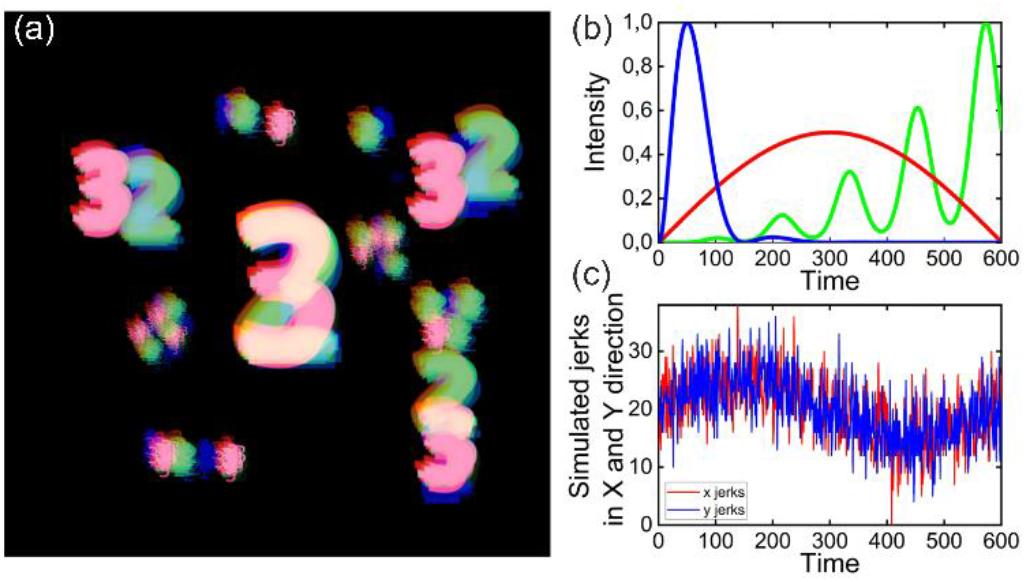
Simulation of a temporal sequence made up 600 images (regular data cube). (a) The time color image distortion corresponding to the average image is displayed. (b) Three different time variations of intensity are indicated by the three colors and correspond to the three numbers displayed on the field of view (same respective colors). (c) The motions (rough simulation of the respiration and heartbeat together with random jerks) graph.

In Fig.10 simulated images before and after the κώ method are displayed using the average time color image distortion. The entropy IQA indicates a value of v = 0.4 as the optimum value for the correction in agreement with the subjective quality from the time color. The image looks sharper but smallest details are not resolved. In the following, we show that this residual blurring is coming principally from the large slow motions characterized by low velocities close to the intensity light variations, fastest jerks being corrected by the κώ method.

**Figure 10.**
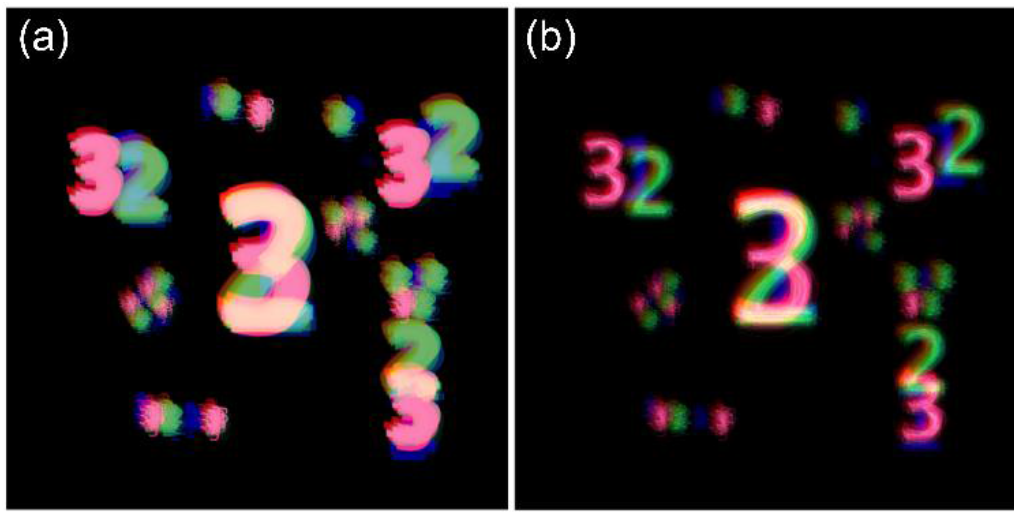
Comparison of the time color image distortion corresponding to the average image before (a) and after (b) the κ ω filter. Entropy IQA indicates a value of v = 0.4 as the optimum value for the correction. Residual blurring is principally coming from large slow motions characterized by frequencies larger than intensity light variations.

In the following, we simulate the jerks as a simple sinusoid defined by A*sin[2πF*t] where A is the amplitude and F the frequency of the sinusoid. We keep the three different time variations of intensity indicated by both the three color and numbers. Fig.11 (a) shows the reference sequence image sharpness case when no motions are simulated is displayed. Fig.11 (b) shows the subfield indicated by a red square for the purpose of a close-up comparison.

**Figure 11.**
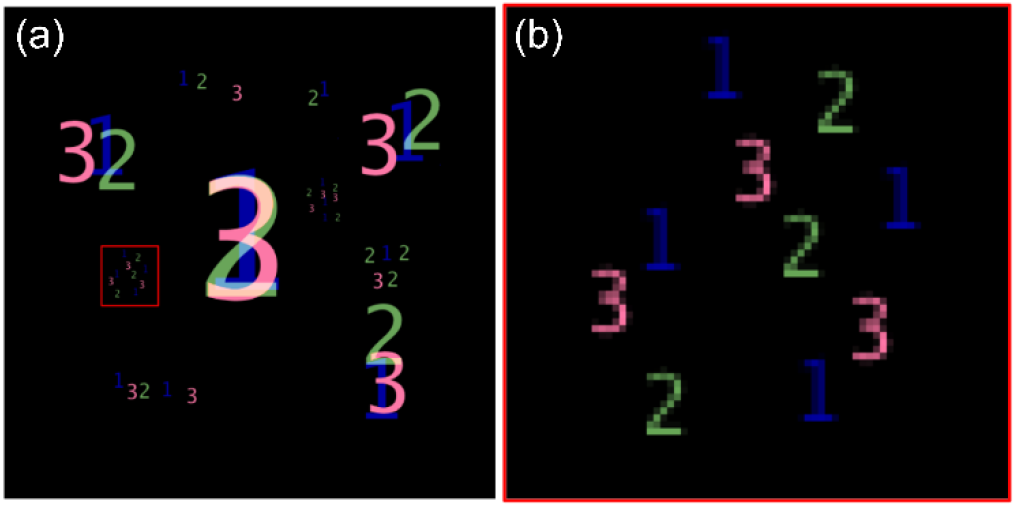
(a) Time color image distortion of the sequence and no motions as a reference. (b) A subfield indicated as red square on the left image and displayed on a growing scale for a close-up comparison.

Fig.12 presents a comparison of the time color image distortion before (a,c) and after the κώ filter (b,d) for two simulated sinusoidal motions A*sin[2πF*t]. It was found that the κώ method is first affected by the frequencies and at a second order by the amplitude. For instance, the processing performs identically for motion amplitudes smallest than 15 pixels when the frequency of the oscillation is 1/300. Entropy IQA indicates the same optimum value for the κώ filter velocity (a value of v = 2). By comparison, in the case of a smaller frequency with a smaller amplitude, more difficult is to compensate the motion closer such as the oscillation with A=5 and F=1/50. The discrepancy in the Fourier space between motions and the variations of intensity is more difficult to determine. Fig. 13 shows the corresponding subfield close-up for the two simulated cases. In this simulation, the sinusoidal frequencies refer to the collection duration time. Longer is the temporal sequence; more efficient is the κώ method.

**Figure 12.**
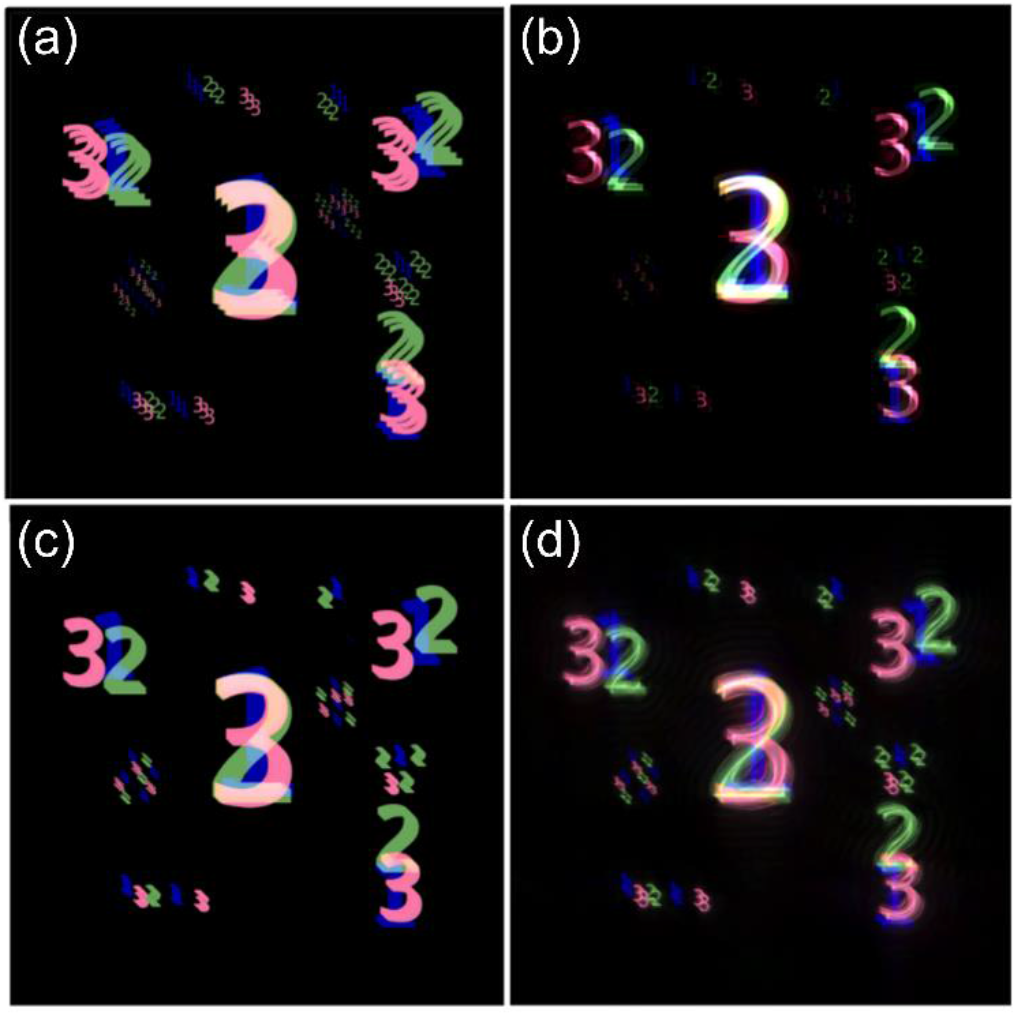
Comparison of the time color image distortion before (a,c) and after the κώ filter (b,d) for simulated sinusoidal motions A*sin [2πF*t]. (a,b) A=10 and F=1/300. (c,d) A=5 and F=1/50.

**Figure 13.**
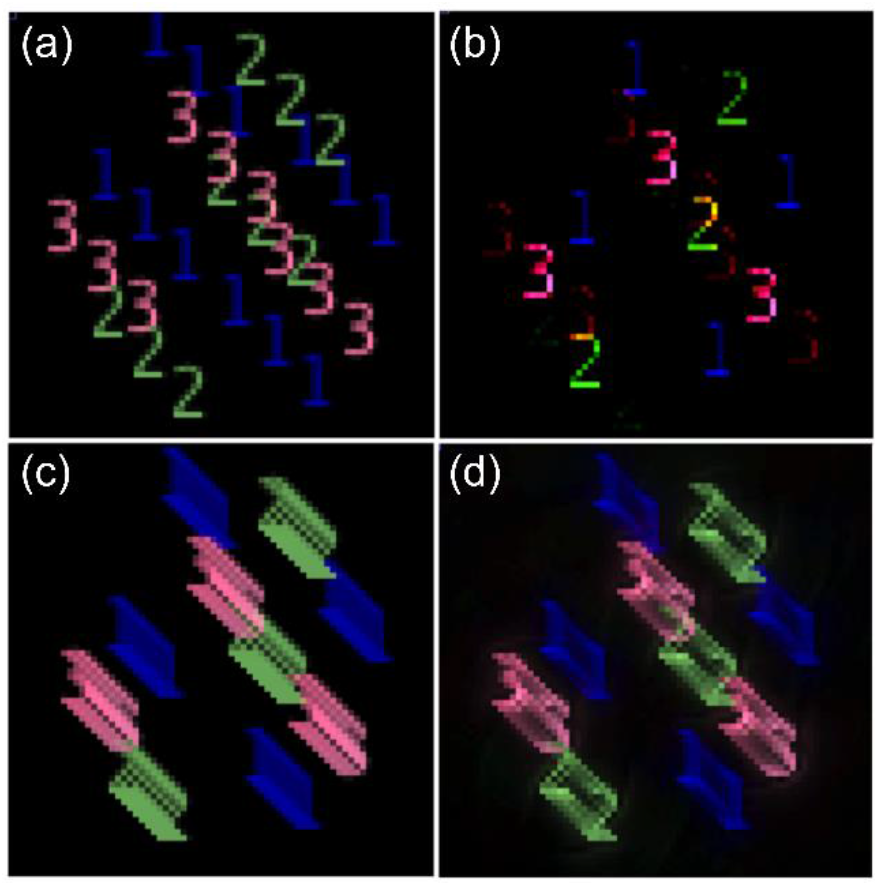
Close-up comparison of the time color image distortion before (a,c) and after the κώ filter (b,d) for simulated sinusoidal motions A*sin[2πF*t]. (a,b) A=10 and F=1/300. (c,d) A=5 and F=1/50. The κώ filter is more efficient for fastest motions in comparison with the intensity variations.

Despite the fact that it is not possible to display a geometric transformation map, the proposed method addresses the problem of removing as many of the distortions as possible while leaving the variations of interest intact. At least, usual registration methods face difficulties to correct occasional images motions when jerks are mixed to the dynamical changes. Image registration technique such as the local image correlation leads to an unsatisfactory result due to the difficulty to determine a static reference all along the temporal sequence. Attempts to determine the relative motion of each image in respect with the previous one in the temporal sequence leads to introduce large saccades that compromise the patterns recognition.

In the case of short temporal sequences or at the end of a fluorescent sequence when the dynamic slows considerably, we applied successfully a rigid registration followed by the κώ filter to improve the spatial resolution. The κώ method is able to preserve the movements of tiny structures of interest, and complete the correction of usual linear registrations.

## 7. Conclusions

We presented a relevant time-space Fourier method so-called κώ method that could be applicable for the stabilization of intra-vital fluorescent in the presence of strong structural changes and significant motion artifacts during acquisition. This method is an alternative to correct variations that are not removed by registration since an exact match is not possible.

Despite fast changes of the structures and intensity with time, this method is able to improve the spatial resolution of the entire sequence, and to reduce the “background noise” created during the fluorescence imaging. Our proposed adaptive approach would be also a useful tool to combine with standard rigid registration or deconvolution methods.

We also proposed a simple approach based on the time visual image distortion method for a quick estimation of the compensation quality. The proposed technique can be used autonomously without supervision and assignation of a reference image, the ideal velocity threshold being determined by testing different velocities and comparing the entropy.

We developed codes both for the Matlab and IDL environments with a similar result although the Fast Fourier Transforms procedures are slightly different. We furnished the Matlab codes of the adapted κώ method depending on the duration of the sequence or/and including a rigid registration step as well as our entropy QIA. The time cost to process a sequence of 600 frames is less than a minute for a current laptop working with Matlab.

## Acknowledgement

The authors would like to thank The Henry Chanoch Krenter Institute for Biomedical imaging and Genomics for the main support of this work (Dr. V. Kalchenko, “Weizmann Staff Scientists Program”), and support provided by COST CA16118 – European Network on Brain Malformations. Professor Meglinski also acknowledge partial support from Academy of Finland (project 326204), MEPhI Academic Excellence Project (Contract No. 02.a03.21.0005) and the National Research Tomsk State University Academic D.I. Mendeleev Fund Program. Special thanks to Dr. K. Miura (EMBL, Heidelberg, Germany) for the Temporal Color Code Image J plugin. Anton Sdobnov also acknowledges the Finnish Cultural Foundation (00180998) grant.

